# Do soil features condition seed germination of gypsum plants? An analysis of the effect of different natural soils

**DOI:** 10.1101/2021.05.13.443982

**Authors:** Nathaniel Heiden, Andreu Cera, Sara Palacio

## Abstract

Gypseous soils are widespread across arid and semiarid environments worldwide. They present remarkable challenges to plants and host a unique flora. We aimed to assess up to what point the specificity and distribution of species on gypsum might be driven by species-specific germination responses to soil gypsum availability. We analyzed the germination of six gypsum specialists and four closely related generalist plant species from the Iberian Peninsula and the Chihuahuan Desert in four different field soils with contrasting concentrations of gypsum, pH and soil texture. Plant restriction to gypsum was unrelated to the germinating ability of seeds on different substrates. Irrespective of their affinity for gypsum, most species germinated better on mixed gypsum-calcareous soil and worse in the acidic soil treatment. Our data suggest soil pH was the main soil feature driving seed germination, while the effect of gypsum content was generally not significant. The main exception was the Iberian gypsum specialist *Helianthemum squamatum* (L.) Dum. Cours., which showed increased germination on gypseous soils and higher germination in response to increased soil gypsum content. Except for this species, our findings indicate alkaline soils favor the germination of most of the species analyzed, irrespectively of their gypsum content.

**Highlights:**

- Gypsum soils are one of the most widespread special substrates of the world
- We evaluated the effects of natural soils on the germination of gypsum plants
- Soil pH was the main soil feature driving seed germination
- The effect of gypsum content was not significant
- We suggest a potential origin of gypsophiles from plants adapted to alkaline soils

## INTRODUCTION

Gypseous soils are a type of alkaline soils containing gypsum (CaSO_4_·2H_2_O) as a main component (Herrero and Porta, 2000). They are spread over 100 million hectares globally, being prevalent in arid and semi-arid regions of the world (Eswaran and Zi-Tong, 1991). Gypseous soils present physical and chemical challenges for plants (Escudero et al., 2015; Verheye and Boyadgiev, 1997). The saturation of the soil solution with Ca^2+^ and sulfate ions (FAO, 1990), suggested to be toxic for some plants (Ernst, 1998), results in alkaline pH, moderate salinity, low nutrient retention capacity (Casby-Horton et al., 2015) and a remarkable nutrient impoverishment of the soil. Gypseous soils typically have hard soil surface crusts, which hamper root penetration and can restrict seedling establishment (Escudero et al., 2000, 1999; Meyer, 1986). They are mechanically unstable due to their lack of plasticity, cohesion and aggregation (Bridges and Burnham, 1980; FAO, 1990); and, in certain areas, they show a low porosity, which limits the penetration of some plant roots (Guerrero-Campo et al., 1999).

The limiting nature of gypseous soils contrasts with their remarkable floristic diversity, rich in endemic and specialized species, which has been defined as a conservation priority of international concern (European Community, 1992; Mota et al., 2011; Ochoterena et al., 2020). According to their specificity to gypsum, plants can be categorized into two general groups: gypsophiles, namely species growing only on gypseous soil; and gypsovags, which grow both on and off gypseous soil (Meyer, 1986). Information on the underlying factors determining gypsophile restriction to gypsum soils is scarce (Escudero et al., 2015). Previous studies have shown how widely distributed gypsophiles from Spain (Palacio et al., 2014, 2007) and the Chihuahuan Desert (Muller et al., 2017) frequently share an ability to accumulate S and Ca in their leaves, which points at a convergent evolution towards similar adaptive traits in plants from these distantly related floras. This pattern could ultimately link to a nutritional specialization of gypsophiles to gypseous soils (Cera et al., 2020). Some gypsum endemic species have also been shown to have a higher ability than co-occurring gypsovags to surpass the hard physical crust typical of gypsum soils at the seedling stage (Romão and Escudero, 2005). The distribution of plants on gypsum soils could also be linked to a differential ability to germinate on gypsum (Romão and Escudero, 2005), since calcium and gypsum concentration can affect seed germination, also in gypsophile species (Anderson, 1982; Cañadas et al., 2014; Merlo et al., 1997; Secor and Farhadnejad, 1978).

Seeds are responsive to their soil environment (Osuna et al., 2015), and germination is one of the riskiest parts of the plant life cycle (Sánchez et al., 2014). Seed germination is largely affected by soil moisture, temperature and light conditions, but soil chemical conditions may also play a role (Baskin and Baskin, 2014). Several studies have described reduced seed germination in response to soil salinity, as a result of negative osmotic potential and ionic toxicity (Pujol et al., 2000; Ungar, 1996 and references therein). High Ca concentrations also decreased the germination of species adapted to acidic soils and favored those of calcicolous species (Anderson, 1982). Soil pH differences also may lead to substantial changes in seed germination (Pierce et al., 1999), and some plant species have been shown to germinate preferentially in a specific pH range (Ma and Liang, 2007). Some widely distributed gypsophile species from Spain and the Chihuahuan Desert increased their germinating ability when treated with a gypsum saturated solution (Cañadas et al., 2014; Merlo et al., 1997; Secor and Farhadnejad, 1978), which some authors attributed to the effect of Ca (Merlo et al., 1997). Seeds of gypsum endemic species may be adapted to germinate in the stressful conditions of these soils, but experimental evidence on the effect of field gypseous soils is lacking (but see Romão and Escudero, 2005).

We analyzed the germinating ability of ten species (Table 1) showing different specificity to gypseous soils on four different natural substrates with contrasting gypsum content, soil texture and pH (Table 2). The species selected include representatives from the two best-known gypsum floras of the world: those of the Iberian Peninsula and the Chihuahuan Desert (Escudero et al., 2015; Moore et al., 2014). Field soils were chosen to cover a broad range in gypsum content and pH and included: gypseous, calcareous, acidic and a 1:1 mixture of gypseous and calcareous soils. We hypothesized that (1) the germination of gypsophiles would be higher in natural gypseous soil and lower in the acidic soil; (2) gypsovags would show similar germination on soil with or without gypsum; and (3) owing to the similar germination responses observed in gypsophiles from the Iberian Peninsula and the Chihuahuan Desert (e.g. Cañadas et al., 2014; Secor and Farhadnejad, 1978), we expect the aforementioned trends would hold true regardless of geographic origin.

**Table 1.** Species included in the study with indication of their taxonomic family, affinity to gypsum soils and geographical origin.

| Species | Family | Gypsum affinity | Origin | Locality | Collection date |
| --- | --- | --- | --- | --- | --- |
| *Gaillardia multiceps*<br>Greene | Asteraceae | Gypsophile | CD | Alkali Lake, New<br>Mexico, US | August 2018 |
| *Helianthemum squamatum*<br>(L.) Dum.Cours. | Cistaceae | Gypsophile | Spain | Villamayor | June 2015 |
| *Helianthemum syriacum*<br>(Jacq.) Dum.Cours. | Cistaceae | Gypsovag | Spain | Villamayor | June 2015 |
| *Lepidium alyssoides* A.<br>Gray | Brassicaceae | Gypsovag | CD | Castillo Formation,<br>Texas, US | August 2018 |
| *Lepidium subulatum* L. | Brassicaceae | Gypsophile | Spain | Villamayor | June 2015 |
| *Matthiola fruticulosa*<br>(Loefl. ex L.) Maire | Brassicaceae | Gypsovag | Spain | Villamayor | July 2015 |
| *Nama canescens*<br>C.L.Hitchc. | Namaceae | Gypsophile | CD | Zaragoza, N.L. Mexico | August 2017 |
| *Nama carnosus* C.L.<br>Hitchc. | Namaceae | Gypsophile | CD | White Sands, Texas,<br>US | September 2018 |
| *Nerisyrenia camporum*<br>(A. Gray) Greene | Brassicaceae | Gypsovag | CD | Socorro, New Mexico,<br>US | August 2018 |
| *Nerisyrenia gracilis* I.M.<br>Johnst. | Brassicaceae | Gypsophile | CD | La trinidad, N.L.<br>Mexico | September 2017 |

CD = Chihuahuan Desert (North America)

**Table 2.**
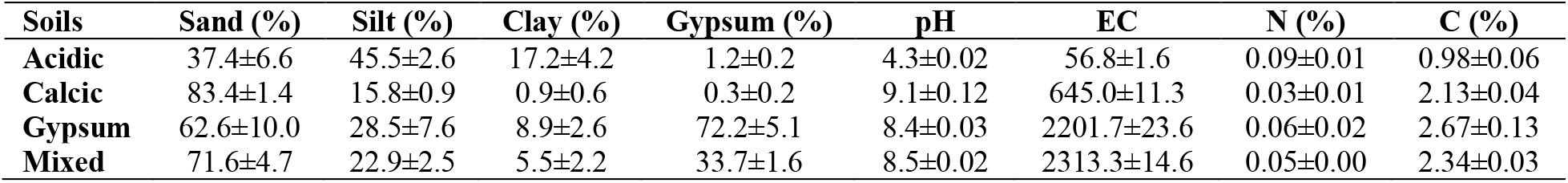
Main physical and chemical features of the different soils included in the experiment. Values are means of *N* = 3 replicates ± SE.

| Soils | Sand (%) | Silt (%) | Clay (%) | Gypsum (%) | pH | EC | N (%) | C (%) |
| --- | --- | --- | --- | --- | --- | --- | --- | --- |
| Acidic | 37.4±6.6 | 45.5±2.6 | 17.2±4.2 | 1.2±0.2 | 4.3±0.02 | 56.8±1.6 | 0.09±0.01 | 0.98±0.06 |
| Calcic | 83.4±1.4 | 15.8±0.9 | 0.9±0.6 | 0.3±0.2 | 9.1±0.12 | 645.0±11.3 | 0.03±0.01 | 2.13±0.04 |
| Gypsum | 62.6±10.0 | 28.5±7.6 | 8.9±2.6 | 72.2±5.1 | 8.4±0.03 | 2201.7±23.6 | 0.06±0.02 | 2.67±0.13 |
| Mixed | 71.6±4.7 | 22.9±2.5 | 5.5±2.2 | 33.7±1.6 | 8.5±0.02 | 2313.3±14.6 | 0.05±0.00 | 2.34±0.03 |

## MATERIALS AND METHODS

### Study species and seed collection

Study species were two Iberian gypsophiles: *Helianthemum squamatum* and *Lepidium subulatum*; two Iberian gypsovags: *Helianthemum syriacum* and *Matthiola fruticulosa*; four gypsophiles native to the Chihuahuan Desert: *Gaillardia multiceps*, *Nama canescens*, *Nama carnosa* and *Nerisyrenia gracilis*; and two Chihuahuan Desert gypsovags: *Nerisyrenia camporum* and *Lepidium alyssoides* (Table 1). Species selection included widely distributed gypsophiles from the two regions of study plus closely related gypsovags for comparison.

Mature fruits were collected from one natural population per species in North America (U.S.A. and Mexico) or Spain. Source individuals were selected to represent the whole population, and included at least 10 different mature individuals separated more than 3 m from each other. Mature fruits from each species were pooled together and, once in the laboratory, they were sorted to collect filled seeds, remove fruit remains and aborted seeds, and then stored at room conditions in paper envelopes from the date of sampling (Table 1) until germination tests were started.

### Soil collection and analyses

Natural gypseous soil was collected from a gypsum outcrop in the Middle Ebro Basin (Villamayor del Gállego, Zaragoza, Spain, 41°41’44.5N, 0°44’26.7”W), calcareous soil was collected from the Iberic System (Ricla, Zaragoza, Spain, 41°30’45.8”N, 1°26’47.8”W), and acidic soil from Moncayo (Agramonte, Zaragoza, Spain, 41°49’40.94”N, 1°49’18.4”W). Soil was collected to 1 m depth by removing O horizons on unfertilized areas in talus slopes and sieved to pass a 1 cm mesh sieve, thoroughly mixed and used to fill pots. A 1:1 mixture of gypseous and calcareous soils was obtained by thoroughly mixing equal volumes of both soils to produce a low gypsum content soil (Table 2).

Three sub-samples were obtained from each soil for physical and chemical analyses. Soil samples were air dried during 2 months at room temperature prior to physical and chemical analyses and subsequently divided in two subsamples, one to be sieved to pass through a 2 mm sieve and the other to remain non-sieved. Sieved soils were used to measure the following variables: gypsum content, measured according to Artieda et al. (2006); soil texture, determined with a particle laser analyser (Mastersizer 2000 Hydro G, Malvern, UK); and soil pH and conductivity, measured with a pH/conductivity meter (Orio StarA215, Thermo Scientific, Waltham-MA, USA) by diluting samples with distilled water to 1:2.5 (w/v) to measure pH and then 1:5 (w/v) to measure conductivity. A subsample of each sieved soil was finely ground using a ball mill (Retsch MM200, Restch GmbH, Haan, Germany) and subsequently used to analyse N and C elemental concentrations with an elemental analyzer (TruSpec CN, LECO, St. Joseph-MI, USA) by EEZ-CSIC Analytical Services.

### Germination experiment

Prior to germination tests, we applied species-specific seed treatments due to the different dormancy-breaking requirements of studied species (Baskin et al., 2006). Seeds from *Helianthemum* species were scarified mechanically with sand paper to break the seed coat that hampers seed re-hydration in these species (Pérez-García and González-Benito, 2006). The seeds of *G. multiceps*, *L. alyssoides*, *N. canescens*, *N. carnosa*, *N. camporum*, and *N. gracilis* were placed in the oven at 50°C for two weeks prior to planting. Previous studies suggested heat shock treatment may increase germination for Chihuahuan desert species (Secor and Farhadnejad, 1978), a positive effect that was confirmed in pre-trials with *N. gracilis* and *N. canescens* (data not shown). No heat scarification is recommended for Iberian gypsophile and gypsovag species (Escudero et al., 1997). Similarly, *L. subulatum* and *M. fruticulosa* were not pre-treated previously, since no previous study indicated the need of dormancy breaking in these species. Seeds from all species were selected under the stereomicroscope to include filled firm seeds with signs of viability. Seeds were disinfected by soaking for 10 minutes in a 5 % bleach solution and then washed with distilled water twice. They were soaked in distilled water for a day before planting. All planting and sterilization were done with autoclaved, ethanol sprayed, and then UV sterilized materials.

Germination trails were set up in two different times in a growth chamber with identical settings, and monitored every two days for 60 days (Cañadas et al., 2014). Between February and April 2019, we analyzed *G. multiceps*, *L. alyssoides*, *L. subulatum*, *N. canescens*, *N. carnosa*, *N. gracilis*, *N. camporum*. Between July and September 2020, we analyzed *H. squamatum*, *H. syriacum* and *M. fruticulosa*. To evaluate seed germination responses to different soil types, 10 replicates of 10 seeds per pot per species were planted in each of the four soil treatments (except for *L. alyssoides* on mixed and acid soil and *H. squamatum* on gypseous soil, which had 9 replicates and *L. subulatum* on acid soil*, N. gracilis* on calcareous soil and *N. camporum* on mixed soil, which had 11 replicates). A total of 400 seeds were planted per species by lightly covering the seeds with soil. Pots of 45 cm^3^ (5 cm × 5 cm × 3 cm) were arranged randomly in a growth chamber with 16 hours of light (flux = 1743-1900 lm, CCT =4000 -6500 K, similar to Escudero et al., 1997) at 25 °C and 8 hours of darkness at 15 °C and 80 % humidity. Alternate 25°C / 15°C temperatures over daily cycles are suitable conditions to promote the germination of Iberian gypsum plants (Escudero et al., 1997; Moruno et al., 2011; Sánchez et al., 2014). Soil surface in pots was kept moist at all times by regularly spraying distilled water. We considered a seed to have germinated once the cotyledons appeared above the soil surface (Wenk and Dawson, 2007). Once counted, emerged seedlings were removed from pots.

### Calculations and statistical analyses

Germination percentage was calculated as the percentage of seeds that germinated within 60 days per pot. Cumulative counts of germinated seeds were used as the response variable in statistical models evaluating the effects on plant germination. Data were not normal and variances increased with mean values, consequently, we used generalized linear mixed models (GLMMs) fixed to a Poisson distribution with log link function. “Affinity to gypsum”, “soil treatment” and “geographical origin” were included as fixed effects, and “species” nested within “taxonomic family” as random effects, to test for differences in germination among plants. Similar GLMMs with “soil treatment” and “geographical origin” as fixed effects and “species” nested within “taxonomic family” as random effects were run separately for gypsovags and gypsophiles to account for soil effects on the germination of each group of gypsum affinity and geographical origin. Generalized linear models (GLMs) with “Soil treatment” as a fixed factor were run for each species separately. Post hoc Tukey tests were run after GLMMs and GLMs to account for differences among soil types.

To unravel the effect of different soil features on the germination ability of seeds we ran a Principal Component Analysis (PCA) with all soil features measured. This served both to explore the correlation between different soil variables and to summarize the multidimensional space of soil properties into two vectors explaining most of the variability of the dataset. Variables were scaled prior to analysis by subtracting the mean and dividing by the standard deviation, to avoid variation due to different units. The first two PCA axes, explaining 98.5 % of the variability (Supplementary Materials, Table S1) were subsequently included as covariates in GLMMs with “germination counts” as the response variable, “affinity to gypsum” and “geographical origin” as fixed effects and “species” nested within “taxonomic family” as random effects. This aimed at exploring the effect of main soil features, geographical origin and affinity to gypsum soils on the germination ability of plants.

All statistical analyses were run in R 3.6.3. Shapiro Wilk normality tests were run with *nortest* (Gross and Ligges, 2015). GLMMs and GLMs were run with *lme4* package version 1.1-15 (Bates et al., 2007). Multiple comparison Tukey tests were run with *multcomp* package (Hothorn et al., 2009). PCAs were run and visualized with *vegan* package version 2.5-6 (Oksanen et al., 2007), and *ggplot2* package (Wickham, 2009), respectively.

## RESULTS

### Gypsophiles and gypsovags showed similar germination but plants germinated better on mixed soils and worse on acidic soils

In general, gypsovags showed higher germination percentages than gypsophiles across substrates (Table 3, Fig. 1). However, the interaction between gypsum affinity and soil type was not significant, which indicates that the effect of soil type on germination did not differ between gypsophiles and gypsovags (Table 3). Plants germinated better on mixed (low gypsum content) soil than calcareous (*z* = 3.03, *P* = 0.005) or acidic (*z* = 5.96, *P* < 0.001) soils. Also, plants germinated less on acidic soils than mixed, calcareous (*z* = 2.85, *P* = 0.022), or gypseous (*z* = 3848, *P* < 0.001) soils. There was not a significant difference in the germination of seeds planted in gypseous and mixed (*z* = 2.25, *P* = 0.110) or calcareous (*z* = 1.05, *P* = 0.718) soils.

**Fig. 1.**
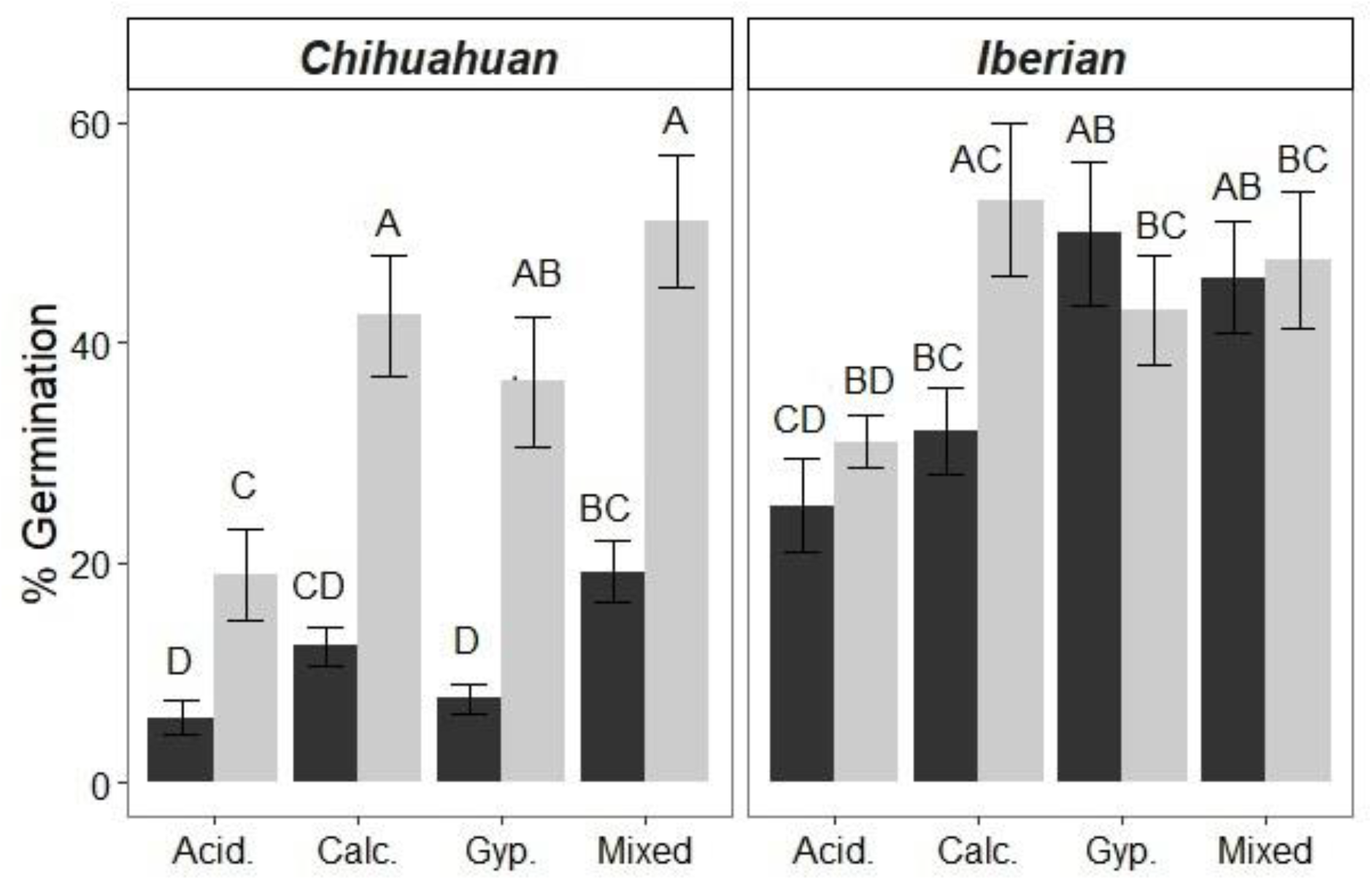
Germination of seeds of gypsophile (black bars) and gypsovag (light grey bars) species from the Iberian Peninsulaand the Chihuahuan Desert sown on different substrates (Acid. = acidic; Calc. = calcareous; Gyp. = gypseous; Mixed = 1:1 gypseous:calcareous soils mixture). Different letters indicate significant differences among soil types within gypsum affinity groups after Tukey post hoc tests (α = 0.05). Values are means ± SE. *N* = 40 for gypsophile and *N* = 20 for gypsovag species from the Chihuahuan desert, except for gypsophiles on calcareous soil and gypsovags on acid soil where *N* = 41 and 19, respectively. *N* = 20 for Iberian species, except for gypsophiles on acid and gypsum soil where *N* = 21 and 19, respectively.

**Table 3.** Results of GLMMs testing: 1) the effect of soil type, plant affinity to gypsum soils (Class), geographical origin (Origin) and their interaction on the seed germination of study species (general model); and 2) the effect of soil type and geographical origin on the seed germination of gypsophiles and gypsovags separately. Models included “Family” and “Species” nested within “Family” as random effects. General model: *N* = 400; Gypsophiles: *N* = 241; Gypsovags: *N* = 159.

|  | General model |  |  | Gypsophiles |  |  | Gypsovags |  |  |
| --- | --- | --- | --- | --- | --- | --- | --- | --- | --- |
|  | *DF* | *Chisq* | *Pr*<br>(*>Chisq*) | *DF* | *Chisq* | *Pr*<br>(*>Chisq*) | *DF* | *Chisq* | *Pr*<br>(*>Chisq*) |
| **Soil Type** | 3 | 64.87 | < 0.001 | 3 | 37.89 | < 0.001 | 3 | 33.53 | <0 .001 |
| **Class** | 1 | 8.88 | 0.003 | - | - | - | - | - | - |
| **Origin** | 1 | 24.38 | < 0.001 | 1 | 22.25 | < 0.001 | 1 | 6.60 | 0.010 |
| **Soil Type:Class** | 3 | 6.49 | 0.090 | - | - | - | - | - | - |
| **Soil Type:Origin** | 3 | 14.20 | 0.003 | 3 | 17.49 | < 0.001 | 3 | 5.21 | 0.157 |
| **Class:Origin** | 1 | 9.62 | 0.002 | - | - | - | - | - | - |
| **Soil_Type:Class:Origin** | 3 | 8.41 | 0.038 | - | - | - | - | - | - |

The analysis of results of each species separately indicated some species-specific responses. For example, the gypsum specialist *H. squamatum* showed a remarkable increase in germination in gypseous soils as compared to other substrates (Fig. 2). Most of the rest of species, either gypsophiles or gypsovags, showed trends consistent with the general response, germinating better in mixed soil and worse in acidic soil, whereas calcareous and gypseous soils showed intermediate germination. Exceptions to this general trend were *G. multiceps* and *H. syriacum*, which showed no significant differences in the germination among substrates.

**Fig. 2.**
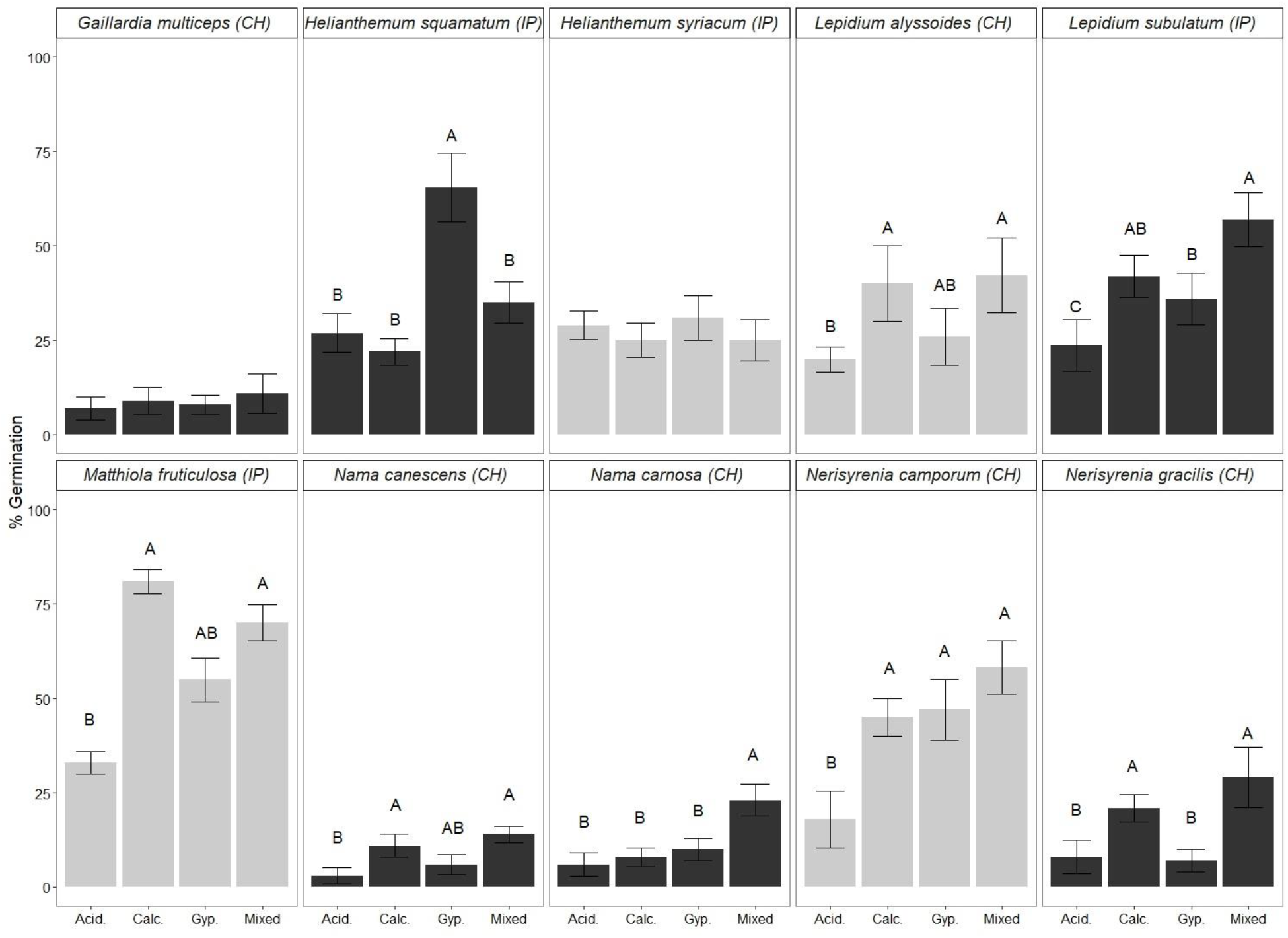
Box plots showing differences in the germination percentage (%) of study species species from the Iberian Peninsula (IP) and the Chihuahuan Desert (CH) cultivated on different substrates (Acid. = acidic; Calc. = calcareous; Gyp. = gypseous; Mixed = 1:1 gypseous:calcareous soils mixture).. Different letters indicate significant differences among soil types after Tukey multiple comparisons tests (α = 0.05). *N* = 10, except for *L. alyssoides* on mixed and acid soil and *H. squamatum* on gypseous soil, where *N* = 9, and *L. subulatum* on acid soil*, N. gracilis* on calcareous soil and *N. camporum* on mixed soil, where *N* = 11.

Seeds from Iberian species germinated generally better than those from Chihuahuan plants (Table 3; Fig. 1), this was partly due to species like *N. canescens, N. carnosa, G. multiceps* or *N. gracilis,* which showed remarkably low germination in general, most values falling below 20 % (Fig. 2). This effect was more important in gypsophiles than gypsovag species. Accordingly, the interaction between gypsum affinity and seed origin was also significant: Iberian gypsophiles showed higher germination percentages than those from Chihuahua (Table 3, Fig. 1). Finally, the interaction between seed origin and soil treatment was significant, since Iberian plants germinated generally better on gypseous soils than Chihuahuan plants. This was particularly due to the effect of *H. squamatum*. Indeed, the analysis of the effect of this interaction separately for gypsum specialists and generalists plants indicated that the interaction was only significant for gypsophiles, highlighting the effect of *H. squamatum* on the high germination of Iberian gypsophiles on gypsum. In accordance to these results, the triple interaction was highly significant (Table 3). Despite these differences due to the generally lower germination in seeds from Chihuahuan desert plants, patterns of variation in response to soil treatments were largely similar among different study species, regardless of the geographical origin, showing a increased germination on alkaline over acidic substrates (Fig. 2).

### Soil pH rather than gypsum content was the main soil feature driving seed germination

PCA of soil features indicated the variability in our soil treatments could be summarized in two main axes, which, together, explained 98.5 % of the variability (Supplementary Materials, Table S1). The first PC was related to soil pH, total C content, EC, total N and soil texture, while the second PC was mainly driven by the gypsum content and, to a lower extent, EC (Supplementary Materials, Table S2). Consequently, the first PC separated acidic soil samples (with lower pH, higher N and a silty-clayish texture) from those of alkaline soils with a higher pH but also higher EC, total C (likely due to the carbonate content) and a sandy texture (Fig. 3). This axis explained two thirds of the variation in the dataset (Supplementary Materials, Table S1). Contrastingly, PC2 segregated alkaline soils owing to their gypsum content and EC, and explained 24.3 % of the variability. The inclusion of these two PCA components as covariates in GLMMs showed only PC1 was significant in the general model (Table 4), highlighting the effect of soil pH, rather than gypsum content and EC, on the germination of study species. The interaction between gypsum affinity of plants and PC2 was significant, indicating the germination of gypsophiles and gypsovags responded differently to gypsum content in the soil. Accordingly, while the effect of PC2 was not significant for gypsovags (Table 4), it became significant when analyzed separately in gypsophiles (Table 4). This trend was largely due to the response of *H. squamatum*, and disappeared when this species was excluded from the analyses (data not shown). The geographical origin of plants also interacted with the effect of PC1 (Table 4), highlighting the overall higher germination percentages of seeds from Iberia, also in the acidic substrate (Fig. 2). Finally, the significant triple interaction between origin, affinity for gypsum and PC2 highlights the differential response of gypsophiles from different origins to soil gypsum content. Whereas the percentage of gypsum in the soil was unrelated to seed germination of Chihuahuan Desert gypsophiles, it significantly affected the germination of Iberian species, mainly through the response of *H. squamatum* (see above).

**Table 4.** Results of GLMM testing: 1) the effect of plant affinity to gypsum soils (Class), the geographical origin of seeds and the two main components (PC1 and PC2) of the PCA of main soil features on the germination of study species (general model); and 2) the effect of soil PCA components 1 and 2 on the germination of gypsophiles and gypsovags separately. “Family” and “Species” nested within “Family” were included as random effects. General model: *N* = 400; Gypsophiles: *N* = 241; Gypsovags: *N* = 159.

|  | General model |  |  | Gypsophiles |  |  | Gypsovags |  |  |
| --- | --- | --- | --- | --- | --- | --- | --- | --- | --- |
|  | *DF* | *Chisq* | *Pr (>Chisq)* | *DF* | *Chisq* | *Pr(>Chisq)* | *DF* | *Chisq* | *Pr(>Chisq)* |
| **Class** | 1 | 9.02 | 0.003 | - | - | - | - | - | - |
| **Origin** | 1 | 25.16 | < 0.001 | 1 | 24.07 | < 0.001 | 1 | 5.89 | 0.015 |
| **PC1** | 1 | 50.80 | < 0.001 | 1 | 20.76 | < 0.001 | 1 | 30.00 | < 0.001 |
| **PC2** | 1 | 1.59 | 0.208 | 1 | 5.64 | 0.018 | 1 | 0.14 | 0.710 |
| **Class:Origin** | 1 | 9.45 | 0.002 | - | - | - | - | - | - |
| **Class:PC1** | 1 | 0.00 | 0.958 | - | - | - | - | - | - |
| **Class:PC2** | 1 | 3.95 | 0.047 | - | - | - | - | - | - |
| **Origin:PC1** | 1 | 4.78 | 0.029 | 1 | 1.85 | 0.174 | 1 | 2.95 | 0.086 |
| **Origin:PC2** | 1 | 0.80 | 0.371 | 1 | 6.02 | 0.014 | 1 | 0.77 | 0.381 |
| **Class:Origin:PC1** | 1 | 0.01 | 0.938 | - | - | - | - | - | - |
| **Class:Origin:PC2** | 1 | 5.91 | 0.015 | - | - | - | - | - | - |

**Fig. 3.**
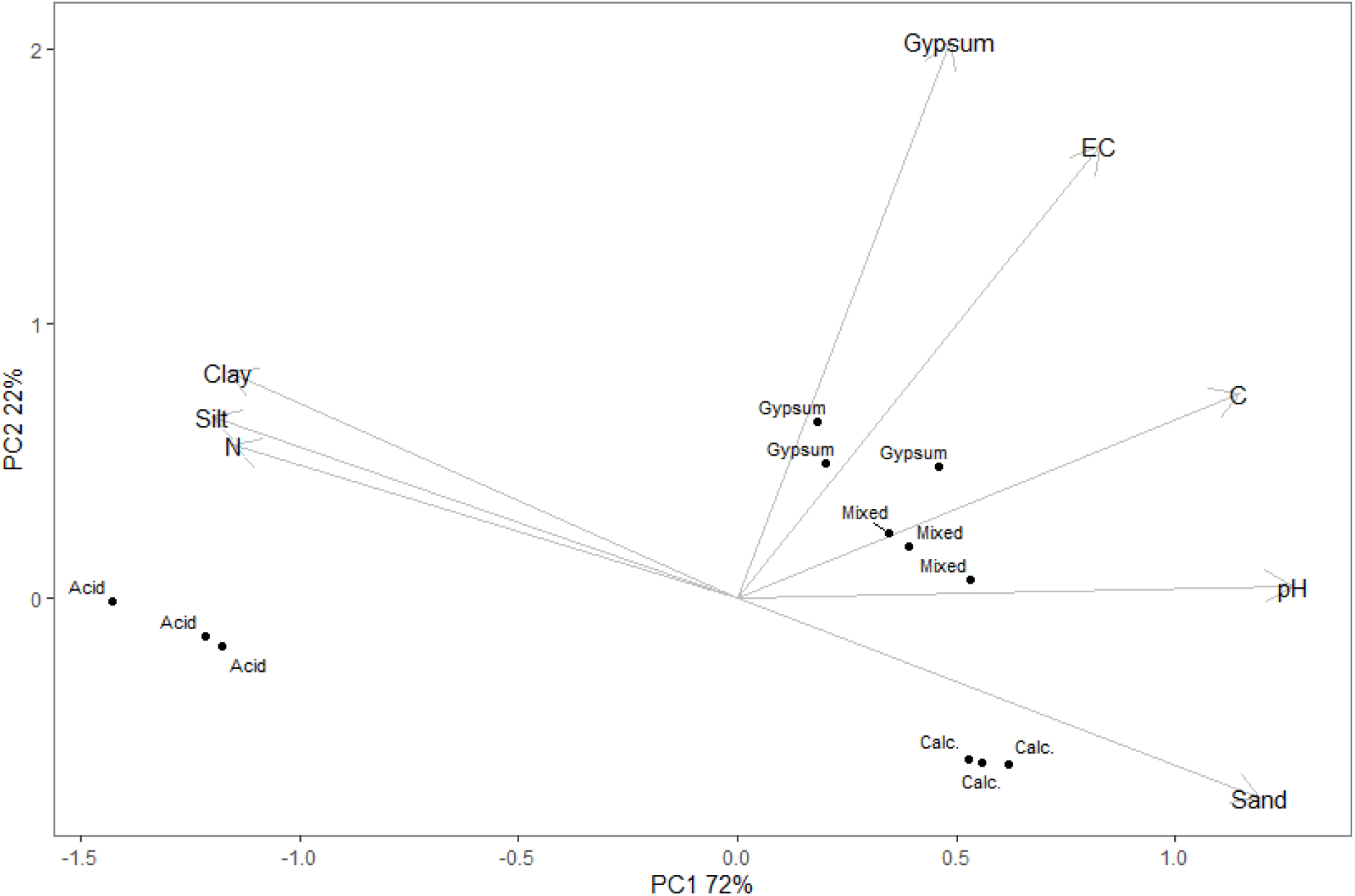
Plot showing the results of PCA of the different soil types included in the experiment and their relation to the main soil features analyzed.

## DISCUSSION

Contrary to our expectations, the restriction of gypsophiles to gypsum observed in the wild did not alter their ability to germinate on a broad range of soils with different chemical and physical features. Similarly to the gypsovags analyzed, they germinated better on alkaline over acidic soils, irrespective of soil gypsum content. These results generally stood independently of seed origin. Most species analyzed showed the highest germination in mixed gypseous:calcareous soil, with a low gypsum content and alkaline pH, whereas germination was decreased in acidic soils. The higher germination of most plants on the mixed soil treatment could be related to the relatively more favorable conditions (such as lower pH) in this soil type as compared to the rest of alkaline soil treatments. These results seem to indicate germination alone does not drive the restriction of gypsophiles to natural gypsum soils.

Exceptions to these general trends were the gypsophile *G. multiceps* and the gypsovag *H. syriacum* which showed no differences in the germination among substrates. The gypsum specialist *H. squamatum* also departed from the general trend, showing increased germination on gypseous soils compared to the rest of substrates. These results are in agreement with the sole previous study analyzing the effect of field soils on the germination of gypsophiles (Romão and Escudero, 2005). In their analysis, Romão and Escudero (2005) evaluated the germination of *Helianthemum squamatum* on natural gypseous and calcareous soil and on a standard nurse substrate. They reported no significant differences on the seedling emergence on different types of substrates, but significantly higher survival of seedlings over 3 months on natural gypsum soils vs. commercial nursery substrate. Taken all together, these results point at a potential specialization of the seeds of *H. squamatum* to germinate and establish on soils with high gypsum content.

Similar to Romão and Excudero (2005), we found that the content of gypsum in the soil was unrelated to the germination ability of most of our study species. However, previous studies with the addition of CaSO_4_ solutions to seeds of gypsophile and gypsovag species showed that increased CaSO_4_ favored the germination of widely distributed gypsophiles (Merlo et al., 1997), including *Lepidium subulatum* (Cañadas et al., 2014) and *Gaillardia multiceps* (Secor and Farhadnejad, 1978), which was taken as evidence of the positive effect of gypsum on the germination of gypsophiles. These discrepancies can be explained by the different experimental approaches in these studies. Instead of chemical solutions, our study and that of Romão and Escudero (2005) used field soils. This approach is closer to natural conditions (Ma et al., 2015), since it entails other physico-chemical factors such as soil texture, soil pH or EC. Although gypsophiles germinated generally better on mixed soils, their germination rates were also high on gypseous soils, in accordance to their occurrence on this type of substrates.

The analysis of the effect of different physico-chemical features of soils on the germination ability of seeds highlighted pH as a driving factor, with seeds of most species being favored by alkaline pH. Plant hormones, such as auxins and ethylene, play an important role in induction of germination, and the levels of these hormones have been shown to be affected by soil pH (Ribeiro et al., 2018). pH may also alter element toxicity, with subsequent effects on germination (Abedi et al., 2013). Most of the species included in this study grow frequently on alkaline soils, either gypseous (all gypsophiles), calcareous (*N. camporum*, *L. alyssoides*, *H. syriacum*) or dolomites (*H. syriacum*). Consequently, it is not surprising that they show better performance on soils with relatively high pH.

While pH was the soil feature showing a higher loading on the PC component explaining most of the variability in the germination of seeds, other factors correlated to pH, such as total C or soil texture, could also play a role. In our alkaline soils, total C may likely be the result of carbonate content, since the amount of organic matter in natural gypseous and calcareous soils is frequently low (Casby-Horton et al., 2015) and carbonates are a frequent component of gypseous soils in northeastern Spain (Palacio et al., 2007). As such, total C in our analysis may be a reflection of the alkaline nature of the soils and, consequently, highly correlated to pH.

In our analysis, soil texture (sand, silt and clay content) was a strong component of PC1, contributing to the separation of alkaline and acidic soils. The acidic soil included in our experiment had higher clay and silt content than alkaline soils, which were sandier. While soil texture is an important factor affecting the water holding capacity of soils and hence seed germination (Wenk and Dawson, 2007), it is unlikely that these factors played a role by themselves in our experiment. First, soils were kept constantly moist at all times throughout the development of the experiment. And second, silt and clay (which may favor soil water retention over sand) had negative loadings in PC1 and hence negatively affected germination. This result can only be explained by the circumstantial correlation between silt and clay and acidic pH derived from the design of the study.

Although not directly accounted for in this study, Ca^2+^ availability is a key factor affecting the germination ability of seeds of many plant species (Anderson, 1982). Merlo et al. (1997) suggested that the main effect of high CaSO_4_ on gypsophile germination was through an increase in Ca^2+^ in the soil solution. Gypseous soils have a higher Ca^2+^ activity than calcareous soils (FAO, 1990). Consequently, we could expect Ca^2+^ availability in our study to correlate with gypsum content in the soil, and hence display a comparatively low effect on the germination of seeds than soil pH. Similarly, previous studies analyzed the separate effect of Ca^2+^ availability and pH on the germination ability of seeds, and concluded that the concentration of calcium compounds in the soil may be of primary importance at the germination stage due to their effects on pH, rather than their nutritional content (Pierce et al., 1999; Rorison, 1960).

Seeds from Iberia germinated generally better than those from the Chihuahuan desert. Pereira et al (2021) recently reported similarly low germination percentages in the American gypsum-associated species *Arctomecon californica*, which they attributed to the need of desert species to experience multi-year conditioning prior to germination, a trait related to the existence of permanent seed banks in perennial species from gypsum ecosystems (Caballero et al., 2003). We cannot rule out the possibility that germination conditions in our experiment might have been slightly cool for Chihuahuan Desert species. Secor and Farhadnejad (1978) germinated seeds at cycles of 23-29 °C, slightly warmer than our 15 – 25 °C. These authors also applied cold stratification to perennial seeds prior to germination, a pre-treatment not included in our experiment. However, despite the differences in overall seed germination, species from Iberia and the Chihuahuan Desert showed similar responses to different soil treatments, highlighting the higher germination of most species on alkaline vs. acidic soils, irrespectively of the gypsum content.

## CONCLUSIONS

Our study did not find support for the hypothesis that plant specificity for gypsum is dependent, in part, on preferential germination in gypseous soils. This was true for most species both from Spain and from North America, except for the gypsum specialist *H. squamatum*, which showed increased germination on gypseous soils compatible with a selection at the germination stage. In the rest of species analyzed, however, we found a common trend among gypsophiles and gypsovags of increased germination in alkaline soils (both calcareous and gypseous) and lower germination in acidic soils. According to this evidence, it appears unlikely that the distribution of most species on gypsum is largely driven by differential germination based on soil type. In contrast, soil variables other than gypsum content may be more important determinants of germination in most of our focal species. Our results are compatible with a potential origin of gypsophile lineages from calcicolous species.

## Acknowledgements

We are grateful to María Pérez-Serrano, Pablo Tejero, José Azorín, Sara Alberdi, Lola Echevarria and Laura de La Puente for help with experiment set up and maintenance; Gabriel Montserrat-Martí, Pedro Sánchez and Ismael González for help with soil collection; Jesús Revilla and Antonio Palma for assistance with growth chamber conditioning; Isabel Llorca, Zöe Feder, Victoria Puértolas, Clara Blasco, Natalia Revilla and Jorge Sereno for help with seed processing; Míriam Aixart for advice on seed biology; Rebecca Drenovsky, Clare Muller and Michael Moore, for help with seed supply and useful comments on earlier versions of this manuscript.

## Funding

Funding was provided by MICINN, Spain, [projects CGL2015-71360-P and PID2019-111159GB-C31)] and the European Union’s Horizon 2020 research and innovation program under grant agreement No [H2020-MSCA-RISE-777803]. NH, AC and SP were funded by a Fulbright Research Grant, a FPI fellowship [MICINN, BES-2016-076455] and a Ramón y Cajal Fellowship [MICINN, RYC-2013-14164], respectively.

## Authors’ contributions

All authors designed the study. AC and SP set up the experiment. NH ran the germination tests. All authors analyzed data and wrote the manuscript.

## Conflict of Interest

The authors declare that they have no conflict of interest.

## Notes

### Competing Interest Statement

The authors have declared no competing interest.

