## Supplementary Data for "Do soil features condition seed germination of gypsum plants? An analysis of the effect of different natural soils"

**Table S1.** Eigenvalues and their contribution to the three PCA components of different soil features.

| | **PC1** | **PC2** | **PC3** |
|---|---|---|---|
| **Eigenvalue** | 5.9381 | 1.9438 | 0.11813 |
| **Proportion Explained** | 0.7423 | 0.2430 | 0.01477 |
| **Cumulative Proportion** | 0.7423 | 0.9852 | 1.00000 |

**Table S2**. Loadings of different soil variables after PCA.

| | PC1 | PC2 | PC3 |
|---|---|---|---|
| **Sand** | 2.5377 | 0.7873 | -0.04975 |
| **Silt** | -2.5585 | -0.7165 | 0.05194 |
| **Clay** | -2.4941 | -0.9162 | 0.04562 |
| **Gypsum** | 0.9825 | -2.4294 | 0.44157 |
| **pH** | 2.6531 | -0.0588 | 0.14163 |
| **EC** | 1.7063 | -1.8998 | -0.73579 |
| **N** | -2.4734 | -0.9676 | -0.08982 |
| **C** | 2.3866 | -1.1419 | 0.25002 |
